# Task-Dependent Representation of Feature Binding within the Same Dimension in Visual Working Memory

**DOI:** 10.1101/2023.12.19.572444

**Authors:** Ruoyi Cao, Leon Y. Deouell

## Abstract

Working memory (WM) serves as a neurocognitive system responsible for temporarily storing and manipulating information when its source has disappeared. Previous investigations into whether features within the same dimension were stored separately or conjoined into objects have yielded conflicting findings. Based on increasing evidence on the adaptivity of the working memory system, we conjectured that the format in which objects from the same dimension are stored in Visual Working Memory (VWM) might be contingent on the specific task demands, and subtle distinctions across experiments may account for the disparities in results. In the current study, we introduced modifications to the paradigm used by Luck and Vogel (1997) and Wheeler and Treisman (2002), where the same paradigm led to different conclusions, to assess whether implicit task requirements could influence the storage format of objects in visual working memory. In two conditions, we manipulated the relevance of conjunction between two colors by varying the proportions of Mis-conjunct probes, a probe type depending on conjunction information for accurate responses. The results showed that in both conditions, performance was primarily determined by the number of features rather than the number of objects, aligning with the results of Wheeler and Treisman. Nevertheless, we observed that Mis-conjunct probes, which require conjunction information, exhibited improved performance when tested more frequently. This suggests that the format of retention in working memory, whether in separate or conjoined form, is influenced by the task demands.

## Introduction

Working memory is a cognitive system responsible for the temporary storage and manipulation of information following the removal of external stimuli. It has been the subject of comprehensive investigation (Fukuda, Vogel, Mayr, & Awh, 2010; Vogel, McCollough, & Machizawa, 2005). However, representation formats of objects within working memory are still controversial. Some studies support the notion that conjoined objects are stored, implying that features are bound together into objects. This perspective, known as the “strong object” view, was initially proposed by Luck and Vogel (1997) based on their findings that increasing the number of features within an object did not impair performance in change-detection tasks. They later (Luria & Vogel, 2011) validated this finding through the examination of electroencephalography (EEG) markers for cognitive load, demonstrating that the amplitude of the Contralateral Delay Activity, known to be correlated with the number of items kept in working memory, corresponded to the number of objects stored, irrespective of the number of features within each object. This phenomenon extended to complex objects featuring color–color conjunctions.

Other studies, however, reached different conclusions. For instance, Wheeler and Treisman (2002), based on experiments with a similar paradigm as Luck and Vogel (1997), proposed the “multiple-resources” view. This was based on the observation that task performance declined as the object contained an increased number of features from the same dimension, while it remained constant when these features were from different dimensions. This result suggests that working memory simultaneously maintains features from different feature dimensions, but features from the same dimension compete for storage space. This view was also supported by other studies using similar (Delvenne & Bruyer, 2004; Jiang, Olson, & Chun, 2000; Parra et al., 2009) and different paradigms (Xu, 2002).

This debate between the strong object model and the multiple-resources model is based on the notion that working memory maintenance adheres to a default format, either in the form of bound items or separate features. This notion was challenged by our recent EEG study (Cao, Pertzov, Gao, Shen, & Deouell, 2021), suggesting that WM retention formats of objects consisting of two features from different dimensions, like color and location, are task-dependent. Here, however, we address the retention formats of objects consisting of multiple features from the same dimension, like two different colors.

The retention of features within the same dimension has been identified as a distinct process with possibly different neuroanatomy compared to features of different dimensions. For instance, Parra, Cubelli, and Della Sala (2011) described a brain-damaged patient who exhibited impaired shape-color binding but retained preserved color-color binding abilities. An fMRI study using a visual search paradigm showed that within-dimension conjunction searches prompted increased activation in the left occipital-temporal cortex. In contrast, cross-dimension conjunction searches elicited greater activation in the bilateral intraparietal sulci (Wei, Müller, Pollmann, & Zhou, 2011).

Thus, the primary aim of the present study is to examine whether the flexibility observed in previous research concerning the Visual Working Memory (VWM) retention of features from different dimensions, can be applied to features derived from the same dimension. In an attempt to reconcile the contradictory outcomes reported in the literature, we employed a delayed (Yes/No) recognition task used by Luck and Vogel (1997) and Wheeler and Treisman (2002), which arrived at contradictory conclusions. In this task, participants were presented with a memory array that had to be compared with a probe displayed after a delay period.

A few subtle differences in the design of the above two studies exist which we conjectured could explain the different findings. In Luck and Vogel (1997), each item presented in the memory array contained features randomly selected from all possible feature values with replacement (e.g., more than one item could be red). Therefore, a non-match probe could include an erroneous conjunction of features that appeared in the memory array. In contrast, in the study by Wheeler and Treisman (2002, Experiments 1 and 2), items on the memory array were generated by selecting from possible feature values without replacement (e.g. only one item could be red). Moreover, a non-matched probe always included a feature that had not been used by any item in the memory array. These subtle differences in the design created different task requirements: in the study by Luck and Vogel (1997), bound objects had to be maintained to detect a non-match probe, while in the study by Wheeler and Treisman (2002), remembering a list of all features in the memory array was sufficient to distinguish between match and non-match probe. In the current study, we examined the assumption that these differences in design account for the distinct results.

To that aim, we directly manipulated the proportion of Mis-conjunct probes, which are probes displaying an incorrect combination of two colors present in the memory array and require conjunction information. We posited that a large proportion of such probes would encourage maintenance of conjoined objects, and the results will thus replicate those of Luck and Vogel (1997), which supported the retention of bound objects. In contrast, a low proportion of Mis-conjunct probes would encourage the maintenance of separate features, and the results would align with the findings from Wheeler and Treisman (2002) emphasizing the retention of separate features from the same dimension. We further investigated whether the task could shape the representation formats by comparing the performance for Mis-conjunct probes, which require conjunction information, across the different task conditions.

## Methods

### Participants

A power analysis, conducted based on the findings from Experiment 2 in Wheeler and Treisman’s (2002) study, indicated that a sample of 50 subjects would be required to achieve 98% statistical power with a significance level set at 0.05. We recruited neurologically normal students through the University online recruitment system until we reached a total of fifty individuals (aged 20-30 years, mean±SD 23.50 ±2.45) who met the inclusion criteria. These criteria required an average response accuracy exceeding 60% and a response accuracy of at least 55% in any of the trial types. Subjects who did not meet these criteria may not have fully comprehended the instructions or may not have been sufficiently attentive during the task. Ethical approval for the study was obtained from the ethics committee of the Faculty of Social Sciences at the Hebrew University of Jerusalem, Israel. Participants were randomly assigned to one of two conditions (25 subjects in each group) and received compensation in the form of either two credits or a monetary payment of 60 NIS for their participation.

### Stimuli

Stimuli were generated with Psychotoolbox-3 in Matlab 2018 (The MathWorks, Inc., Natick, Massachusetts). Three distinct types of memory arrays were employed: Two-Item-Two-Color array (2I-2C, see Figure 1a), Two-Item-Four-Color array (2I-4C, see Figure 1b), and Four-Item-Four-Color array (4I-4C, see Figure 1c).

**Figure 1.**
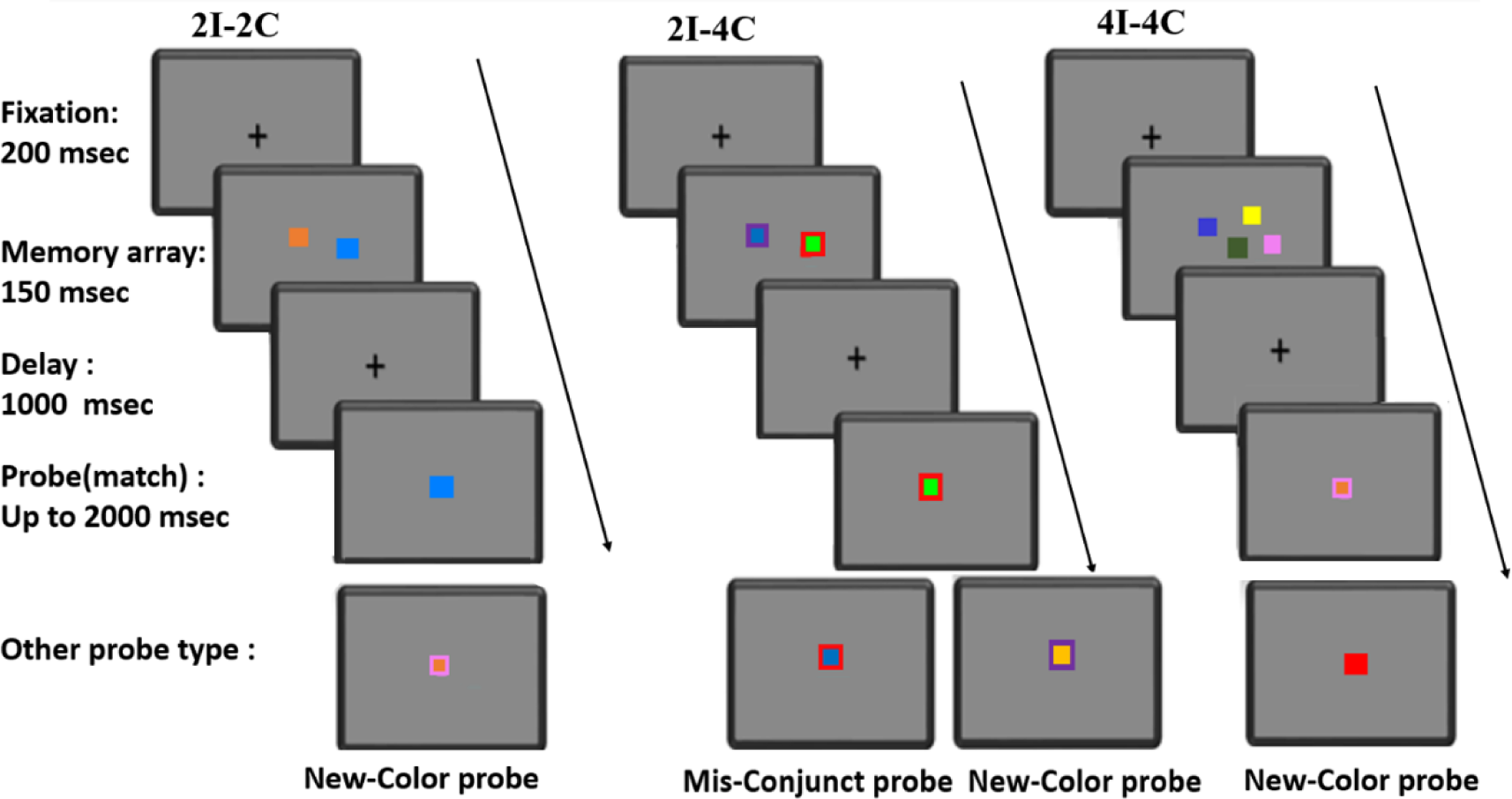
An example of trials with three different types of memory array terminating with a Match probe. Examples of other probe types are presented below each type of trial. The probe types are defined relative to the memory array shown. For example, the pink probe in the 2I-2C is a new-color probe because the array has an orange and blue squares. The same pink probe would have been a match probe in the example 4I-4C trial because it appears in the corresponding memory array. Note also the Mis-conjunct probe which has the inner color of the left item in the array and the outer color of the right item in the array.

Memory arrays consisted of squares that were presented either in a single color or in double colors. The single-color items were displayed in both the 2I-2C and 4I-4C arrays, with each item having a side length equal to 2% of the screen’s width. Since participants used their screens during the experiment, the precise size and visual angle of these items could vary between participants. In the case of the 2I-4C array, double-colored squares were presented with an inner square (side length of 2% of the screen’s width), and an outer frame (side length of 4% of the screen’s width) of a different color. Both the single-color squares and the double-colored squares were randomly positioned within an invisible square (side length: 30% of the screen’s width) centered on the screen.

Colors for the memory arrays were randomly selected without repetition (i.e., each color appeared only once in each array) from a set of nine options (see Figure 1): Dark Green [RGB 0 50 0], Orange Red [255 102 0], Yellow [255 255 0], Medium Purple [128 61 199], Blue [0 0 250], Orange Red [255 102 0], Dodger Blue [0 128 255], Lime [0 255 0], Red [255 0 0], and Light Coral [255 128 153]. Memory arrays containing two similar colors (e.g., blue and Dodger Blue; Red and Orange Red; Light Coral and Medium Purple), or overlapping items were excluded to prevent confusion. This resulted in 120 unique arrays for each of the three types of memory arrays.

Each probe contained only one item presented at the center of the screen, consisting of either a single-colored square or a double-colored square according to the type of memory array it followed. There were three types of probes (Figure 1):

#### Match probes

When following the 2I-2C and 4I-4C arrays (both of which including only single-color squares), a Match probe was a single-color square presented at the center of the screen with a color the same as one of the squares presented in the memory array. When following the 2I-4C array (in which double-color items were presented), a Match probe was a double-color square that had both colors from one of the squares presented in the memory array, with the same conjunction.

#### New-Color probes

When following the 2I-2C and 4I-4C array, a New-color probe was a single-color square presented at the center of the screen with a color that was not present in the memory array of that trial. When following the 2I-4C array, a New-Color probe was a double-color square in which either the inner part color or the outer part color was not present in the previous memory array.

#### Mis-Conjunct probes

This type of probe only followed the 2I-4C array. It was a double-colored item in which both colors were present in the memory array but in an incorrect combination. Specifically, the outer color of one square was conjoined with the inner color of the other square.

### Procedure

Participants downloaded the computer program created using E-prime Go (https://pstnet.com/eprime-go/) and performed the experiment on their computers. There were two conditions: Binding (B) and Feature (F), into which participants were randomly assigned. Each condition started with a practice block. After the practice (18 trials), a cycle of one Induction block (60 trials) followed by two Testing blocks (120 trials each block) was run twice, leading to 600 trials in total (Figure 2). The order of trials in a given block was randomized within each subject.

**Figure 2.**
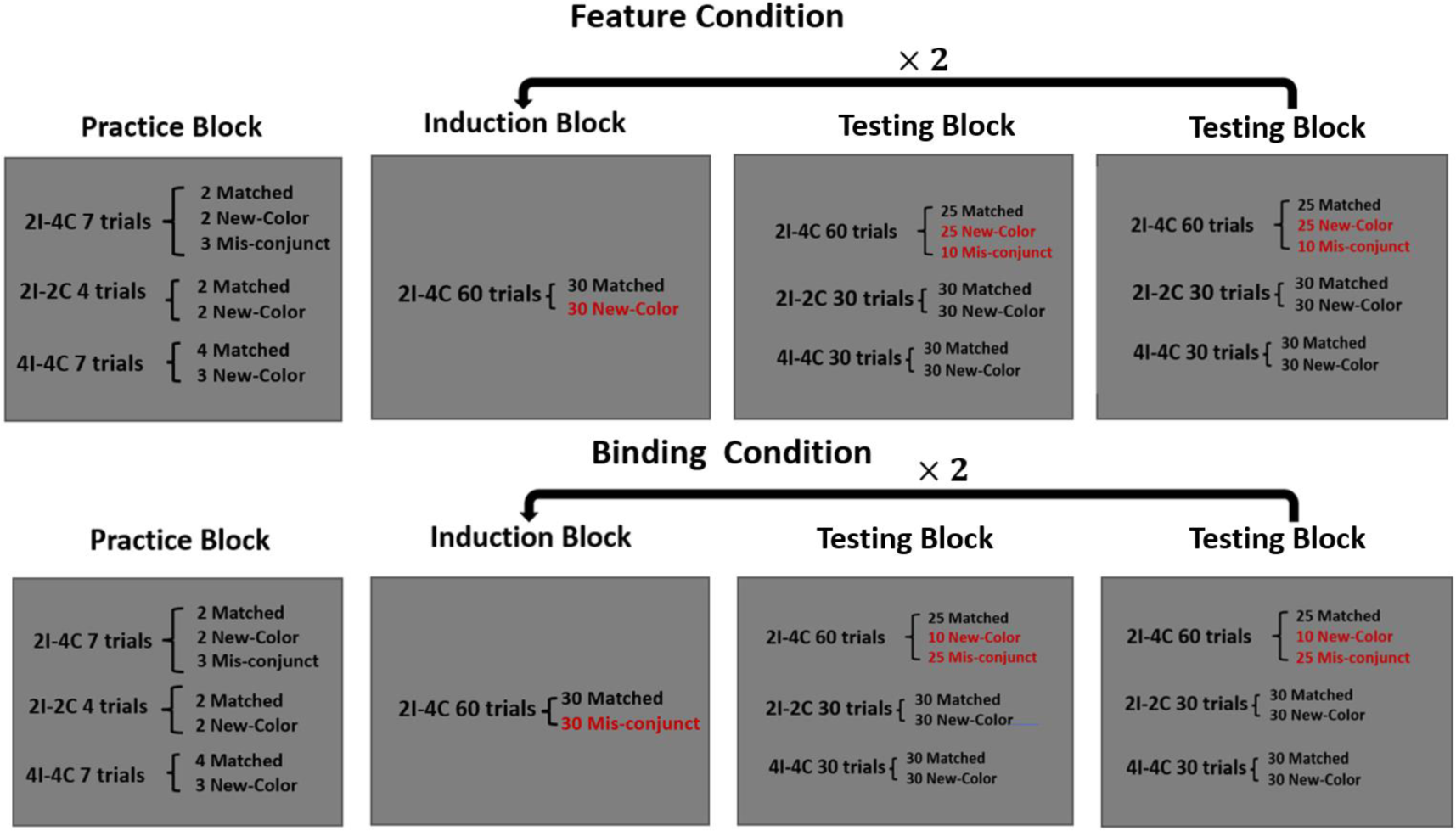
Experiment procedure for Feature and Binding conditions. The difference between conditions (marked in red) is in the type of probes used in the induction block and the proportion of New-color and Mis-conjunct probes in the 2I-4C trials.

#### 1. Instruction and Practice block

Both conditions started with instructions, where subjects learned to respond to all types of memory arrays and probes. Subjects were first shown an example of all possible colors. Then they were instructed to remember items presented on the memory array and to indicate whether the probe presented after a delay period matched one of the items on the memory array or not. They were instructed to indicate match with key “J” and Non-match with key “F”. An example of correct response for each potential probe type following each type of memory array was presented (altogether seven examples). Subjects had to view all examples and to confirm that they understand the rules of the task before continuing to the practice trials. Following these instructions, subjects performed 18 practice trials (See Figure 2 for the number of trials of each type). The order of trials was randomized. As shown in Figure 1, each practice trial began with a 200-millisecond fixation cross presented at the center of the screen. Next, the memory array appeared for 150 milliseconds, followed by a 1-second blank-screen delay period. After the delay period, a probe item was presented, and participants had to press a key to indicate whether the probe item was “Match” (i.e. present in the memory array) or “ Non-match.” The probe disappeared when the response was made or 2 seconds elapsed. Finally, feedback was shown for 300 milliseconds, indicating whether the response was correct, incorrect, or too slow.

#### 2. Induction Block

The objective of the Induction block was to establish a specific maintenance format before the testing blocks. This was achieved by utilizing New-Color probes in the F condition or Mis-conjunct probes in the B condition. Accurate responses to New-Color probes did not necessitate the retention of color conjunction on the same object. In contrast, the retention of conjunction was critical for responding correctly to Mis-Conjunct probes. The trial structure for the Induction Block was the same as that of the practice block. The induction block contained a total of 60 trials with 2I-4C memory arrays, presented in an order randomized for each subject. Among these, half consisted of Match probes (30 trials), while the remaining half comprised of only New-Color probes in the F condition and only Mis-conjunct probes in the B condition. Thus, the type of probes presented in the Induction block was assumed to induce an optimal retention format.

#### 3. Testing Block

Each testing block comprised a total of 120 trials including thirty 2I-2C arrays (half followed by Match probes and half by New-color probes), thirty 4I-4C arrays (half followed by Match probes and half by New-color probes), and sixty 2I-4C arrays. The proportion of Mis-conjunct and New -Color probes following the 2I-4C arrays made the difference in Testing Block between the B and F conditions: in the F condition, 10 trials included a Mis-Conjunct probe, and 25 trials included a New-Color probe. Conversely, in the B condition, 10 trials included a New-Color probe, and 25 trials included a Mis-Conjunct probe. In both conditions, 25 trials included Match probes. Both instruction and the structure of trials in testing blocks were similar to that of the practice blocks, with the exception that no feedback was provided after the response.

### Analysis

Statistical analysis was conducted with JASP (Version 0.17.1; https://jasp-stats.org/) and figures were made by the Python graphic library Seaborn (https://www.python-graph-gallery.com/seaborn/). Only responses in trials of the Testing Blocks were used for analysis.

Response accuracy for each type of probe following each type of memory array was entered into a Condition (B vs F) ×Array type (2I-2C; 2I-4C; 4I-4C) 2 ×3 Mix-measures two-way ANOVA. Degrees of freedom were adjusted for violations of the assumption of sphericity with the Greenhouse–Geisser correction when necessary. In addition, the Inclusion Bayes factor from Bayesian ANOVA test was also calculated. The inclusion Bayes factor is the ratio of posterior inclusion probability and the prior inclusion probability, representing the evidence in the data for including a predictor (van Doorn et al., 2020).

If the variations in task requirements indeed account for the differing results observed in prior studies and the maintenance format is solely influenced by these task demands, we expected to observe a significant interaction between the Condition and Array Type. More specifically, in F condition, we anticipated that the number of features would determine the performance. Thus, we hypothesized that superior performance would be observed in trials involving a 2I-2C array (comprising only two colors) compared to the other two types (both including four colors), which were expected to exhibit similar performance levels. In the B condition, we anticipated that the number of items, rather than the number of colors, would determine the performance. Therefore, performance would be poorer in the 4I-4C array (consisting of four items) compared to the other two types, where only two items were involved (2I-4C and 2I-2C array), while the performances between these two types of arrays (2I-4C and 2I-2C) was expected to be similar. If only a main effect of Array type were found without an interaction, differences in task requirements cannot sufficiently explain the disparities between the findings in the two aforementioned studies. We would proceed to conduct a post hoc analysis with Bonferroni correction, which aimed to determine whether it is the number of features or the number of conjunct items that predominantly influences performance for both conditions.

Additionally, response accuracy for each type of probe following each type of memory array was entered into a 2 ×7 mixed effects two-way ANOVA with factors Condition (between subjects; B vs F) and Probe type (within subjects; Match probe and New Color probe following 2I-2C; Match probe, New Color probe and Mis-conjunct probe following 2I-4C; Match probe and New-Color probe following 4I-4C). The Inclusion Bayes factor from the Bayesian ANOVA test was also calculated. We hypothesized that if indeed the task affects the maintenance format, this effect would manifest particularly for the Mis-Conjunct probe, a probe type that required the retention of conjunctions. In this case, we predicted a significant interaction between Condition and Probe type with performance differences between B and F condition being larger in the Mis-conjunct probe trials than all other probe types combined, which was tested with planed contrast.

By design, response accuracy was emphasized over response time by allowing a relatively long response time (up to 2 seconds). Nevertheless, the above analysis for response accuracy was also applied to reaction time for trials with correct response to confirm that the results from response accuracy were not caused by the time accuracy trade-off.

## Results

Two groups of subjects performed a delayed recognition task in which they decided if a probe item belonged to a previously presented array of items or not (Figure 1). The arrays included two single color squares (2I-2C), four single color squares (4I-4C), or two double-color squares (2I-4C). In critical trials, following the 2I-4C array, the combination of colors was swapped, requiring the subjects to hold the correct conjunction in working memory in order to correctly identify these items as not belonging to the array. Arrays and probes were mixed within blocks. The two groups of subjects differed in the relative proportion of these critical trials requiring memory of conjunction of colors. In the Feature (F) condition, these were relatively rare, thus emphasizing retention of individual colors, whereas in the Binding (B) condition, they were relatively frequent, thus emphasizing the retention of conjunctions of colors. The two groups further went through an induction phase emphasizing either features (in the F group) or conjunctions (in the B group).

Our first hypothesis was that the different emphasis on retaining individual features (colors) or their conjunction will affect the accuracy. Specifically, we hypothesized that subjects in the F condition will be more accurate than subjects in the B condition with arrays requiring only feature memory (the 2I-2C and especially the 4I-4C), and vice versa with arrays requiring conjunction memory (2I-4C), resulting in an interaction between Condition and Array Type. Such an interaction would suggest that previous contradicting results regarding the maintenance format (as features or conjoined objects) may have resulted from differences in experimental design implicitly emphasizing features or conjunctions.

A 2 ×3 Condition (B vs F) ×Array type (2I-2C; 2I-4C; 4I-4C) mixed-measures two-way ANOVA revealed only a significant main effect for Array type, *F(2,96) =255.11, p< .001, η²= 0.56, BFIncl =5.00×10^+36^* (Figure 3a). Post hoc comparisons with Bonferroni correction showed that performance was better with the 2I-2C array than the 2I-4C array, *Mdiff=0.20, SE =0.01, t(49)= 21.49, p< .001*, and 4I-4C array, *Mdiff=0.16, SE =0.01, t(49)= 16.77, p< .001*. Additionally, 2I-4C trials were performed even worse than 4I-4C trials, *Mdiff=-0.04, SE =0.01, t(49)= −4.73, p< .001*. This pattern of results suggested a strong effect of the number of features on VWM performance, with no benefit for combining two features into an object, consistent with the findings of Wheeler and Treisman (2002, Experiments 1 and 2).

**Figure 3.**
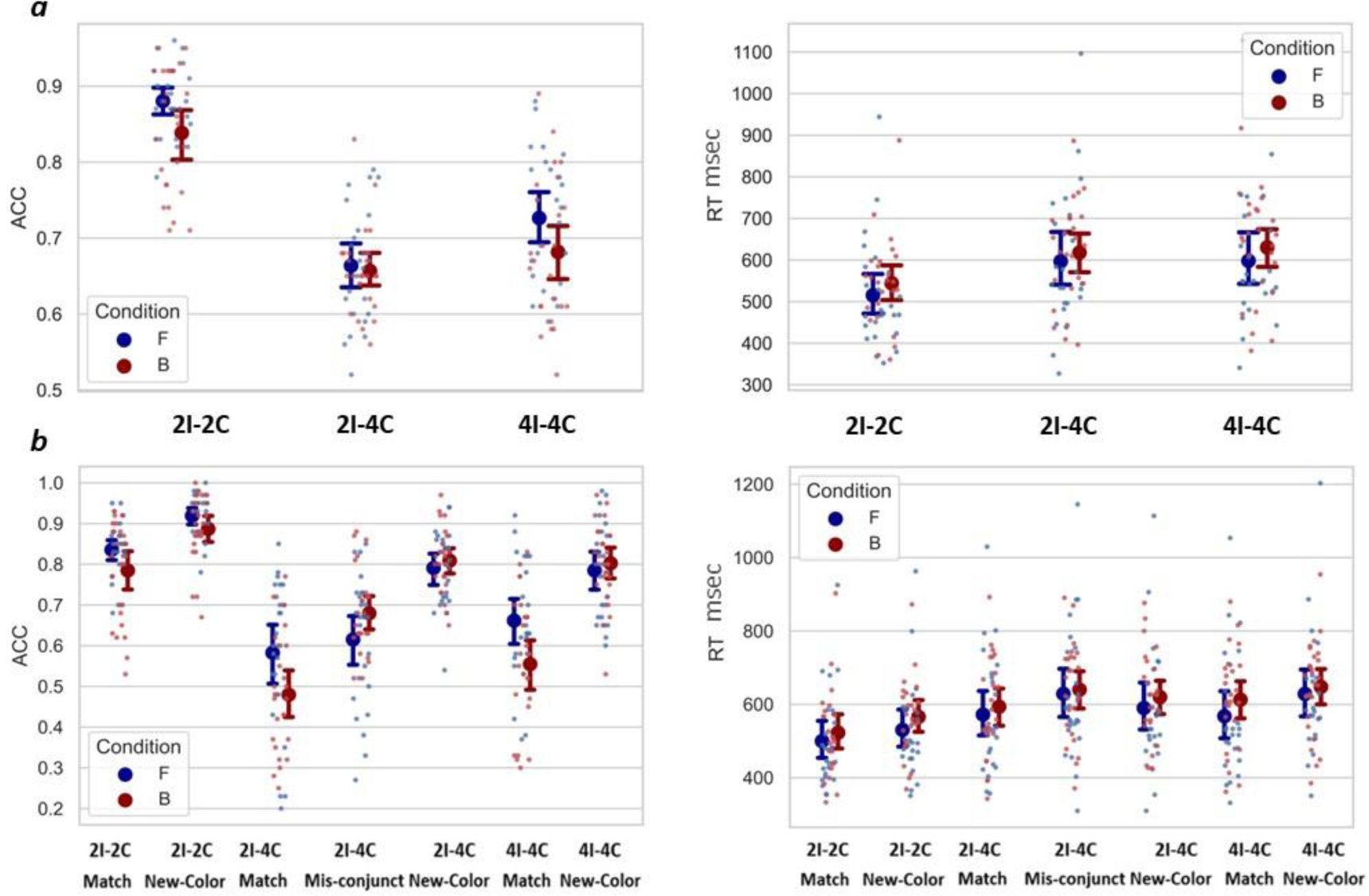
(***a***) Response accuracy (left) and reaction times (right) by array type and condition (***b***) same as (a), but separated by probe type. Small dots represent individual subject means; large dots and error bars depict across subjects mean and 95% confidence interval.

There was a significant performance advantage for the F condition in trials with 2I-2C and 4I-4C arrays, whereas in the 2I-4C case, where conjunctions were relevant, the two conditions resulted in comparable accuracy. Whereas this was in line with our hypothesis that making conjunctions more frequent will improve the retention of conjunctions relative to individual features, the ANOVA indicated only a trend towards an interaction between Array Type and Condition (Figure 3a), *F(2,96) =2.61, p = .08, BFIncl = 0.81*. The main effect of Condition was also not significant, *F(1,48) =2.82, p = .10, BFIncl = 1.01*.

The same ANOVA performed with RT as dependent variable also revealed only a significant main effect for Array type (Figure 3b), *F(2,96) =98.18, p< .001, η²= 0.07, BFIncl =8.242×10^+20^*. We found neither a significant interaction between Condition and Array type, *F(2,96) =0.37, p = .69, BFIncl = 0.17*, nor a significant main effect of Condition, *F(1,48) =0.51, p = .48, BFIncl = 0.66*. Post hoc comparisons for the main effect of Array type with Bonferroni correction suggested that 2I-2C arrays were performed faster than the 2I-4C arrays, *Mdiff=-77.66, SE =6.70, t(49)= −11.60, p< .001*, and the 4I-4C arrays, *Mdiff=-84.43, SE =6.70, t(49)= −12.61, p< .001*. Additionally, we found no significant differences between the 2I-4C array and the 4I-4C, *Mdiff=-6.77, SE =6.70, t(49)= −1.01, p= .31*.

These results aligned with the accuracy findings, indicating that the number of features, rather than the number of items, predominantly influenced the performance. The preceding results did not support the hypothesis that the contradictory findings in the literature resulted from differences in task design. However, it did not discount the potential influence of the task on shaping object maintenance formats. We posited that the impact of the task might be more pronounced with Mis-Conjunct probes, which necessitated conjunction information for accurate responses, as compared to other probe types that did not.

A 2 ×7 two-way mixed-measures ANOVA with factors of Condition (B vs F) and Probe type (Match probe and New Color probe following 2I-2C and following 4I-4C arrays; Match probe, New Color probe and Mis-conjunct probe following 2I-4C) revealed a significant interaction between Condition and Probe type, *F(6, 288) = 4.27, p < .001, η² = 0.03, BFIncl = 70.92 (Figure 3b)*. A main effect of Probe type was also observed, *F(6, 288) = 72.28, p < .001, η² = 0.5, BFIncl = 2.062×10^+51^*, but not for Condition, *F(1, 48) = 2.27, p = 0.14, BFIncl = 0.46*. The planned contrast showed that the performance difference between B and F conditions was greater for the Mis-conjunct probe than other probe types combined, *t(288) = 3.17, p = 0.002*. In fact, performance was significantly better in the B condition than in the F condition in Mis-conjunct probe trials whereas for trials with other probe types the difference was either insignificant or in the other direction. Consistent results were obtained when considering only probes following 2I-4C array (Supplementary S1).

This result suggested that the task demand affect the maintenance format. However, it is important to note that the Mis-Conjunct probes underwent more training and testing in the B condition than in the F condition. Consequently, better performance for these probes in the B condition, compared to other probe types, might be attributed to a practice effect. If this practice effect indeed led to enhanced performance for the Mis-Conjunct probe in the B condition, we should anticipate a mirror result in the F conditions: New-Color probes following the 2I-4C array, which were more frequent in this condition compared to the B condition, would exhibit superior performance compared with all other probe types. Our analysis did not reveal such differences*, t(288) = 1.57, p = 0.12*, suggesting that the advantages of the Mis-Conjunct probe in the B condition may indeed be a genuine effect of the task requirement emphasizing conjunctions.

The speed-accuracy trade-off did not account for the observed effects in response accuracy. A Condition ×Array type 2 ×7 Mix-measures two-way ANOVA revealed a main effect of Probe type (Figure 3b), *F(6, 288) = 48.15, p < .001, η² = 0.08, BFIncl =5.26×10^+37^*. Importantly, we found no significant interaction effect between Condition and Probe type, *F(6, 288) = 0.72, p = 0.63, BFIncl = 0.042*, nor a main effect of Condition, F*(1, 48) = 0.47, p = .50, BFIncl =0.66*.

## Discussion

We probed visual working memory (VWM) in conditions that either encouraged or discouraged the retention of conjunctions between two colors within an item. Our study revealed a consistent pattern: the number of colors, rather than the number of conjoined items, predominantly influenced the performance. This suggests that features originating from the same dimension are primarily retained separately within VWM. However, the results also suggests that the maintenance format is not completely rigid. Conjunction information was better preserved when it had greater task relevance, even implicitly. This provides evidence for flexibility in working memory retention formats for objects composed of features from the same dimension.

In the experiment, we manipulated the relative proportion of Mis-Conjunct probes and New-Color probes to create conditions where conjunctions between two colors within an item were task-relevant to different degrees. Across both conditions, our results consistently indicated a significantly lower performance for four colors compared with two colors, regardless of the number of items. This outcome aligned with the observations made in Wheeler and Treisman’s studies (2002, Experiments 1 and 2), as well as with the results from other studies attempting to replicate their findings.

Olson and Jiang (2002) conducted a series of experiments involving the manipulation of color saturation, the use of blocked or mixed conditions, variation in the duration of memory images, and the introduction of verbal load. All these manipulations led to the same outcome: the performance in working memory tasks was predominantly influenced by the number of colors rather than the number of objects. Another study conducted by Jing, Chengbo, and Qiang (2018) proposed that two colors might be integrated into meaningful objects but not into meaningless objects. However, their findings demonstrated that regardless of the meaningfulness of the objects, colors from the same objects were not integrated. The current study explored a novel factor, the implicit task relevance, and nevertheless provided evidence in favor of the above findings.

While the differences in task design were insufficient to account for the contradictory results of Wheeler and Treisman (2002) and Luck and Vogel (1997), the manipulation did affect performance. Performance for Mis-Conjunct probes was better when conjunction information was more frequently required by the task. This finding added to previous literature on the flexibility of working memory retention. Previous studies showed that VWM can selectively maintain task-relevant visual features while ignoring irrelevant ones that were simultaneously presented (Griffin & Nobre, 2003; Park, Sy, Hong, & Tong, 2017; Pertzov, Bays, Joseph, & Husain, 2013; Ye, Hu, Ristaniemi, Gendron, & Liu, 2016). The present study went beyond the task relevance of individual features, showing that the VWM can also be adjusted based on task-relevance of conjunction information. Moreover, such flexibility was also found between features from the same dimension.

The manipulation of the task relevance of conjunction between features from the same dimension, with different types of probes, was tested in one previous study (Parra, Cubelli, & della Sala, 2011). Similar to our study, they found a significantly lower accuracy in the “Conjoined Color Condition” compared to the “Independent Color Condition”, researching the same conclusion suggesting features in the same dimension are retained separately in VWM. However, the design did not allow to disentangle the effects of task relevance on encoding or retention from reaction to the probe, since both were varied between conditions. In our study, by varying the proportion of the different probes, we were able to reveal the effect of task relevance by making comparison within the same probe type.

It is worth noticing that our study used an indirect way to estimate formats of representation in working memory. Like the study of Wheeler and Triesman and that of Luck and Vogel, it followed the reasoning that working memory performance inversely depends on the number of units stored (Luck & Vogel, 2013). Since there are presumably only a limited number of “slots” available at any moment, working memory performance indicates the number of slots being occupied, allowing inference about formats of representation in working memory. However, the results of the current study, showing worse performance in the 2I-4C condition than the 4I-4C condition, could be interpreted in another way: even if features from the same dimension have been combined into a single storage unit, such units consume more memory resources than those that store a single feature, resulting in decrease of performance. Gajewski and Brockmole (2006) distinguished between the object-unit hypothesis and the independent-stores hypothesis with an approach that involved examining the recall frequencies of both individual features and combinations of features. They posited that under the object-unit hypothesis, features from the same object would be recalled or forgotten together based on an all-or-none principle, whereas the independent-stores hypothesis proposed the opposite. This approach, as well as many other potential paradigms without assuming the slot model, should be applied for future studies seeking to understand the formats of object during visual short-term memory maintenance. Additionally, the current study applied only one time duration, limiting the inference on whether the task manipulated the initial encoding, the later retention or the finally retrieval process. Further investigation is needed to clarify these issues.

To conclude, our study applied the paradigm used by Luck and Vogel (1997) and Wheeler and Treisman (2002) to test whether the task requirement could manipulate the results and thus reconcile previous contradictory findings. However, we found that despite the task relevance, number of features, rather than number of item, determined performance, which aligns with the results from Wheeler and Treisman. Meanwhile, by demonstrating a better performance on probes requiring conjunction information when tested more frequently, our study provided new evidence that the memory of conjunction between features from the same dimension is task-dependent. This emphasizes the importance of considering the impact of task requirements, even implicit ones, when conceptualizing and studying the binding problem in working memory.

## Acknowledgement

We thank Neta Licht for discussion about design and analysis while helping with running experiments.

## Notes

### Competing Interest Statement

Leon Y. Deouell is a co-founder, shareholder and consultant of Innereye Ltd., a neurotech start-up company. However, the activities of Innereye Ltd. are not related to the research presented in this manuscript.

### Summary of Updates

1: The title of the paper was updated from "Task-Dependent Representation of Feature within the Same Dimension in Visual Working Memory" to "Task-Dependent Representation of Feature Binding within the Same Dimension in Visual Working Memory" 2: Mis-conjunction probe was replaced by Mis-conjunct probe consistently.

