## Supplement S1 for "Task-Dependent Representation of Feature Binding within the Same Dimension in Visual Working Memory"

Supplementary **S1**

A 2 × 3 two-way mixed-measures ANOVA with the factors of Condition (B vs F) and Probe type (New Color probe & Mis-conjunct probe following 2I-4C) on accuracy revealed a significant interaction between Condition and Probe type, *F(2, 96) = 4.77, p = .011, η² = 0.04, BFIncl =9.70* . A main effect of Probe type was also observed, *F(2, 96) = 46.25, p < .001, η² = 0.40, BFIncl = 7.06×10^+14^*, but no main effect was found for Condition, *F(1, 48) = 0.13, p = 0.72, BFIncl = 0.21*. The Planned contrast showed that the performance difference between B and F conditions was greater for the Mis-conjunct probe than other probe types combined, *t(96) = 2.25, p = 0.03*. Planned contrasted showed that performance differences between F and B was not significantly greater in F than B for New-Color probes than the other two probe types, *t(96) = 0.77, p =0.44,* suggesting that practice effect might not account for the task effect found on Mis-conjunct probes.

A 2 × 3 two-way mixed-measures ANOVA with the factors of Condition (B vs F) and Probe type (New Color probe & Mis-conjunct probe followed 2I-4C) on RT revealed a significant main effect of Probe type, *F(2, 96) = 15.54, p < .001, η² = 0.02, BFIncl =12404.83*, but neither the significant main effect for Condition, *F(1, 48) = 0.26, p = 0.61, BFIncl = 0.64*, nor the interaction, *F(2, 96) = 0.49, p =0.61, BFIncl = 0.16* . With no interaction between Probe type and Condition, the result from response accuracy is not likely due to trade-off between time and the accuracy.
